# Systematic identification of A-to-I editing associated regulators from multiple human cancers

**DOI:** 10.1101/812610

**Authors:** Tongjun Gu, Audrey Qiuyan Fu, Michael J. Bolt, Xiwu Zhao

## Abstract

A-to-I editing is the most common editing type in human that is catalyzed by ADAR family members (ADARs), ADAR1 and ADAR2. Millions of A-to-I editing sites have been discovered recently, however, the regulation mechanisms of the RNA editing process are still not clear. Here we developed a two-step logistic regression model to identify genes that are potentially involved in RNA editing process in four human cancers. The first step by classifying the editing sites into different categories assists the analysis at the second step. In the first step, ADAR1 was identified as the enzyme that associated with the majority of the A-to-I editing sites. Thus, ADAR1 was taken as a control gene in the second step to identify genes that have a stronger effect on editing sites than ADAR1. In addition, the detectable interferons and their receptors were used as covariates in the both steps to account for potential association caused by interferons. Using our advanced method, we successfully found a set of genes that were significantly positively or negatively associated (PA or NA) with specific sets of RNA editing sites. We highlighted two genes, *SRSF5* and *MIR22HG* which were supported by multiple evidences. Most PA and NA genes were unique to each cancer, and only a few shared across two cancers. Pathway enrichment analysis showed that the PA genes from the four cancer types were enriched in Immune System, while the NA genes were enriched in two pathways: Metabolism of RNA, and Metabolism. The functional similarity of the PA and NA genes across all the four cancers indicates that even though most of the editing associated genes were unique to each cancer, they may impact on editing process through common pathways. Interestingly, the PA genes from kidney cancer were enriched for survival-associated genes while the NA genes were depleted of these genes, indicating that the PA genes may play more important roles in kidney cancer development.

## Introduction

RNA editing is one of the post-transcriptional mechanisms that can diversify the transcriptome by changing the sequences of RNA, providing the flexibility for organisms to adapt themselves to different internal and external environment. The most common type of editing in human is adenosine to inosine (A-to-I) and inosine can be read as guanine by reverse transcription and translation machineries. A-to-I editing is catalyzed by ADAR gene family, which contains three major members, ADAR1, ADAR2 and ADAR3, however, the function of ADAR3 is still not clear^1^. Both ADAR1 and ADAR2 are synthetic lethal genes when either of them was knocked out in mouse models. Homozygous deletion of ADAR1 in mice can lead to death before E12.5 with severe defects in primitive and definitive hematopoiesis^2^. Although mice with homozygous deletion of ADAR2 can survive after birth, the mice die within 20 days with defects in motor neuron degeneration and motor function abnormalities^3^. ADAR proteins must dimerize to be functional^4,5^. They can form either homodimer or heterodimer, consequently, the relative abundance of ADAR1 and ADAR2 is important for RNA editing biogenesis and function^4^. In recent years, millions of RNA editing sites have been identified, but the regulation of RNA editing process and cofactors of the enzymes remain largely unknown.

Recently it was demonstrated that RNA editing plays a leading role in the development of hepatocellular carcinoma^6^. They found a causal RNA editing site, then further determined the corresponding enzyme. They found that the status of the editing efficiency matches the expression of the corresponding enzyme in the populations they studied^7^, indicating that the underlying genetics can affect both editing efficiency and gene expression. In our previous and several other groups’ studies^8-10^, a strong genetic effect on RNA editing was identified. Here we developed a two-step generalized linear regression model to detect the novel genes potentially involved in the RNA editing process based on the association of gene expression with editing sites using hundreds of cancer samples. The first step was to identify which enzyme the editing sites were significantly associated with. The second step was to identify the genes that have potentially stronger regulation effect on RNA editing sites than the primary enzyme: the genes were significantly associated with editing sites while the primary enzyme became not. And we found a set of high-confidence candidates that were associated with specific sets of RNA editing sites.

We found that the candidate regulators were not common to all editing sites and tissue specific. But some cancers shared a large portion of same candidate genes, such as ∼45% of the genes positive associated (PA) with editing sites in LUAD overlapped with the genes in KIRC although only 11 LUAD PA genes were found. Pathway enrichment analysis showed that the function of the PA genes was enriched in Immune System pathway. And the function of negatively associated (NA) genes was enriched in Metabolism of RNA and Metabolism pathways. The results indicate that although most genes were unique to each cancer, they most likely work through similar pathways. Interestingly, the KIRC PA genes were significantly enriched with clinical associated genes while KIRC NA genes were not. Both the pathway analysis and the survival data support that the PA and NA genes may play different roles in cancer development.

## Results

### The Process for the identification of RNA editing associated genes

We quantified a total of 3,800 editing sites, which were discovered from our previous study^11^, in four human cancers with a total of 1,635 RNA-seq samples: liver hepatocellular carcinoma (LIHC), lung adenocarcinoma (LUAD), kidney renal clear cell carcinoma (KIRC), and thyroid carcinoma (THCA). The sample information, including sample size and sequencing depth, is listed in Supplementary Table 1.

We developed a two-step logistic regression model with the goal to identify high-confidence RNA editing associated genes. In the model, the binomial distribution was used to estimate RNA editing ratio by taking the number of edited reads as the number of successful events. The sequencing depth is not even for each editing site within and across samples. Because the logistic regression model considers the sequencing depth at each site, it was more accurate and sensitive than using the empirical ratio (calculated as the number of edited reads / total number of reads at each site). The first step was to classify the editing sites into different categories based on the association with each enzyme. Three enzymes, ADAR1, ADAR2 and ADAR3, were involved in RNA editing process, however, in our samples, only ADAR1 and ADAR2 were expressed at detectable level (mean expression > 1 RPKM (Reads Per Kilobase of transcript per Million mapped reads)) (Figure 1a). Thus, ADAR1 and ADAR2 were tested in the first step. We found that in all the four cancers, ADAR1 was the dominant enzyme significantly associated with the majority of the editing sites (The highest number of ADAR2 associated sites in KIRC was still ∼13 fold less than the number of ADAR1 associated editing sites in KIRC, Table 1). Therefore, we selected the editing sites that were significantly associated with ADAR1 for the second step regression. In the second step, the statistical model was the same as the first step except a gene was added as a covariate. Thus, in additional to test the association between ADAR1 and the editing site, the association between the gene and the same editing site was examined. Two criterions were used to find the editing associated genes. First, the gene was significantly associated with the editing site. Second, ADAR1 switched from an associated gene in the first step to an unassociated gene with the editing site in the second step. The two criterions indicate that the gene can explain the variance of the editing site across samples more than ADAR1, and thus the gene was potentially a stronger regulator in RNA editing process than ADAR1. In our model, ADAR1 was taken as a control gene in the process of detecting editing sites associated genes. The detectable interferons and interferon receptors, gender and race were included as covariates in both steps to exclude their potential effects. Figure 1b showed the logic of the two-step logistic regression. The details of the model are in the methods section.

**Table 1.** The number of editing sites in the ten categories calculated based on the association of RNA editing sites with ADARs in KIRC, LUAD, THCA and LIHC.

| Categories | KIRC | THCA | LUAD | LIHC |
| --- | --- | --- | --- | --- |
| ADAR1+/ADAR2+ | 12 | 2 | 7 | 1 |
| ADAR1+/ADAR2# | 570 | 467 | 1542 | 623 |
| ADAR1+/ADAR2- | 390 | 706 | 96 | 237 |
| ADAR1#/ADAR2+ | 49 | 13 | 8 | 2 |
| ADAR1#/ADAR2# | 1748 | 1564 | 1453 | 2170 |
| ADAR1#/ADAR2- | 91 | 34 | 6 | 4 |
| ADAR1-/ADAR2+ | 14 | 3 | 3 | 3 |
| ADAR1-/ADAR2# | 6 | 3 | 1 | 1 |
| ADAR1-/ADAR2- | 1 | 1 | 0 | 5 |
| ADAR1%/ADAR2% | 856 | 899 | 584 | 522 |

Notes: +, significantly positive association with p value less than 1.3e-05 (FDR at ~5% after Bonferroni correction); -, significantly negative association with p value less than 1.3e-05; #, not significantly association with p value higher than 1.3e-03; %, un-classified. For example, the first category, ADAR1+/ADAR2+, contains the editing sites that were significantly positive associated with both ADAR1 and ADAR2. The categories, ADAR1+/ADAR2+, ADAR1+/ADAR2# and ADAR1+/ADAR2-, were highlighted in light blue and were selected for the second step of logistic regression.

**Figure 1.**
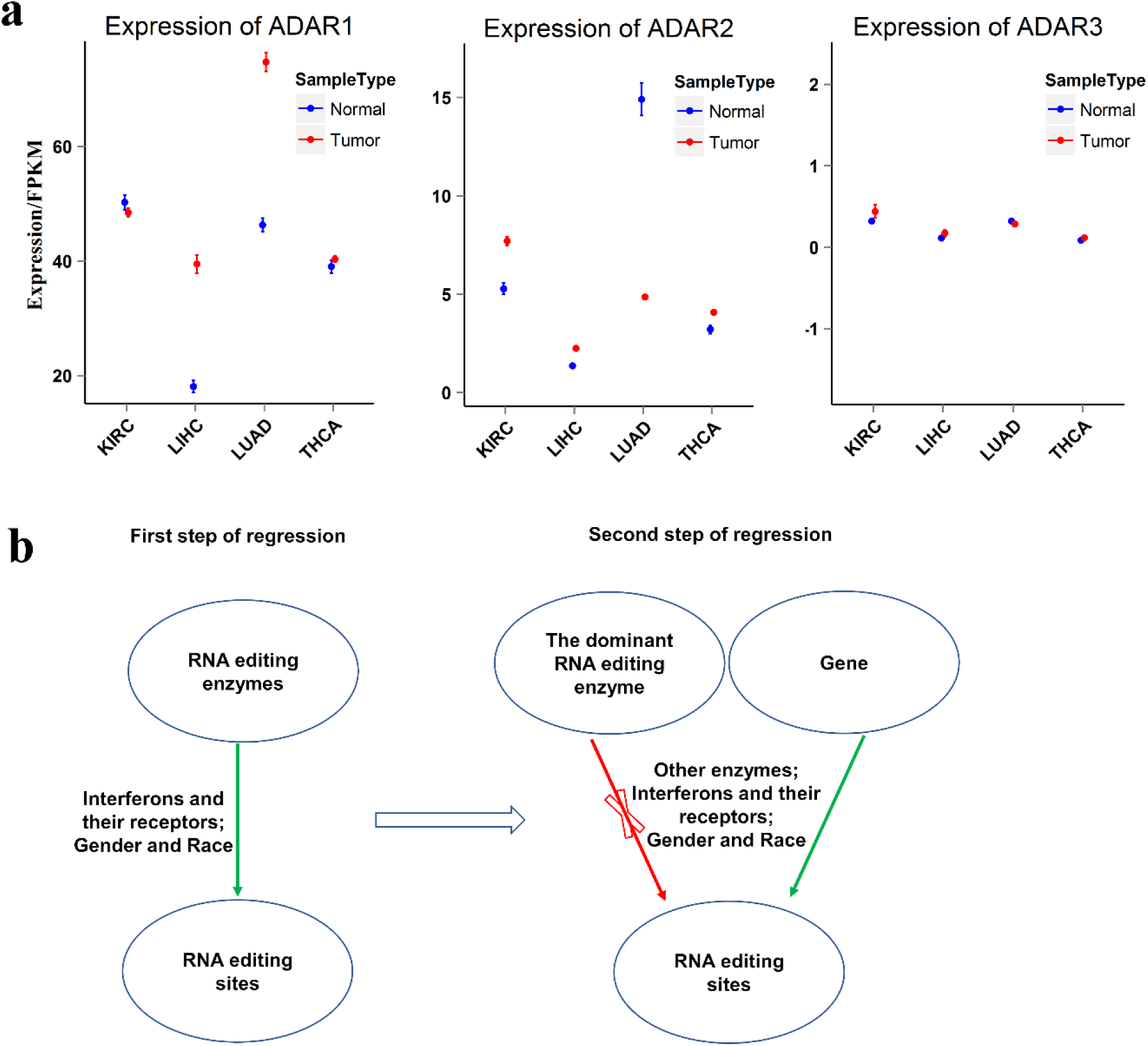
The expression of ADARs in the four cancers and the model of the two-step logistic regression. (a) The expression of ADAR1, ADAR2 and ADAR3 in KIRC, LIHC, LUAD and THCA. FPKM: Fragments Per Kilobase of transcript per Million mapped reads. (b) The model used for discovering genes that associated with RNA editing sites. The first step was to identify the dominant ADAR that catalyzed RNA editing sites, which was ADAR1 in the four cancers. Therefore, we chose the editing sites that were significantly associated with ADAR1 for the second step analysis. The second step was to find the genes that involved in RNA editing process by choosing the genes that can better explain the variance of RNA editing efficiency than the ADARs: the genes were significantly associated with RNA editing sites while the enzyme became not associated in the second step. Because ADAR1 was the enzyme that associated with the majority of the editing sites in all four cancers, we chose ADAR1 as a control gene in the second regression step. The covariates used in the regression were: other enzymes (ADAR2 in our study), interferons and their receptors, gender and race.

### Identify enzyme-specific sets of A-to-I editing sites from the first step of logistic regression

To best clarify the association between RNA editing sites and the enzymes, we grouped the editing sites into 10 categories based on the association with ADARs (Table 1): ADAR1+/ADAR2+, ADAR1+/ADAR2#, ADAR1+/ADAR2-, ADAR1#/ADAR2+, ADAR1#/ADAR2#, ADAR1#/ADAR2-, ADAR1-/ADAR2+, ADAR1-/ADAR2#, ADAR1-/ADAR2- and ADAR1%/ADAR2%. The + and - represent the editing sites were significantly positive (+) or negative (-) associated with the enzyme at threshold of FDR 5% (p value ∼1.3e-05) after Bonferroni multiple correction. # represents the editing sites were not significantly associated with the enzyme at threshold of p value > 1.3e-03. % represents the editing sites were in between +/- and #. The number of editing sites in each category is listed in Table 1. Here we used a stringent multiple testing correction method, Bonferroni multiple correction, to find the strong RNA editing sites-associated ADARs^12^. Therefore, the sites in ADAR1# or ADAR2# or ADAR1%ADAR2% related categories were still possible associated with ADARs, but not as strong as the sites in ADAR1+ or - and ADAR2+ or - related categories.

Although most editing sites (category ADAR1#/ADAR2# and ADAR1%/ADAR2%) did not pass the high threshold to be called as sites that significantly associated with either ADAR1 or ADAR2, there were hundreds of editing sites in ADAR1+/ADAR2# that significantly positive associated with ADAR1 (Table 1). In contrast, only a few sites (the highest number was 49 in KIRC) were in ADAR1#/ADAR2+. The significant difference between the two categories may be caused by the difference between the expression of ADAR1 and ADAR2. The expression of ADAR1 was much higher than ADAR2 (Figure 1a), indicating ADAR1 plays a major role in editing in the four cancer types. There were a set of editing sites significantly positive associated with both ADAR1 and ADAR2 (the highest number was 12 in KIRC). Previous studies showed that in *Drosophila* ADARs can form functional heterodimer^4^. However, while no evidence of functional heterodimerization has been found in human, maybe these sites were edited by ADAR1 and ADAR2 separately in a manner that some transcripts were edited by ADAR1 while some were edited by ADAR2.

We collected a few RNA editing sites from several studies to validate our analysis results. Three editing sites have been tested with both computational and experimental work in human liver tumor samples: editing sites in *AZIN1, FLNB* and *COPA*^6,7^. *The authors found that AZIN1* editing was catalyzed by ADAR1^6^, *COPA* was catalyzed by ADAR2^7^, and *FLNB* can be catalyzed by both^7^. We obtained consistent results: *AZIN1* editing was associated with the expression of ADAR1 and *COPA* editing was associated with the expression of ADAR2 in all the four cancer types (Figure 2a). In our populations, *FLNB* editing was only significantly associated with ADAR1 (Table 2). Another editing site, the editing site in *FLNA*, was reported to be catalyzed by ADAR2 in mouse brain samples^13^. We showed here that *FLNA* editing was also significantly associated with ADAR2 (Table 2). The consistency of our results with previous studies on known editing sites indicates the high precision of our method, and also indicates that the catalytic activity of the enzymes was conserved across tissues and species.

**Table 2.** Examples of the association for several RNA editing sites with ADARs in KIRC, LUAD, THCA and LIHC. *AZIN1* and *FLNB* editing were significantly positive associated with ADAR1 while *COPA* and *FLNA* editing were significantly positive associated with ADAR2.

|  |  | KIRC |  | LUAD |  | LIHC |  | THCA |  |
| --- | --- | --- | --- | --- | --- | --- | --- | --- | --- |
|  |  | Pval | Coef | Pval | Coef | Pval | Coef | Pval | Coef |
| AZIN1 | ADAR1 | 4.17E-109 | 0.014 | 0 | 0.011 | 0 | 0.026 | 2.67E-195 | 0.031 |
| AZIN1 | ADAR2 | 2.36E-47 | -0.039 | 0.015 | -0.0075 | 4.63E-15 | -0.11 | 1.14E-23 | -0.075 |
| FLNB | ADAR1 | 1.28E-113 | 0.013 | 0 | 0.010 | 1.14E-215 | 0.020 | 3.18E-303 | 0.031 |
| FLNB | ADAR2 | 1.51E-18 | -0.020 | 9.89E-06 | -0.015 | 1.14E-11 | -0.091 | 1.42E-54 | -0.086 |
| COPA | ADAR1 | 1.30E-47 | -0.0090 | 2.48E-11 | -0.0025 | 8.58E-10 | -0.0083 | 4.51E-10 | -0.011 |
| COPA | ADAR2 | 5.80E-265 | 0.081 | 3.68E-197 | 0.16 | 2.37E-48 | 0.33 | 2.67E-104 | 0.20 |
| FLNA | ADAR1 | 1.49E-54 | -0.0052 | 5.73E-143 | -0.0057 | 2.59E-18 | -0.012 | 5.79E-112 | -0.017 |
| FLNA | ADAR2 | 0 | 0.092 | 0 | 0.16 | 5.14E-30 | 0.16 | 0 | 0.23 |

Notes: Pval is the p value calculated from the first step regression. Coef is the coefficient calculated from the first step regression. Positive number in Coef represents positive association when Pval is smaller than 1.3e-05.

**Figure 2.**
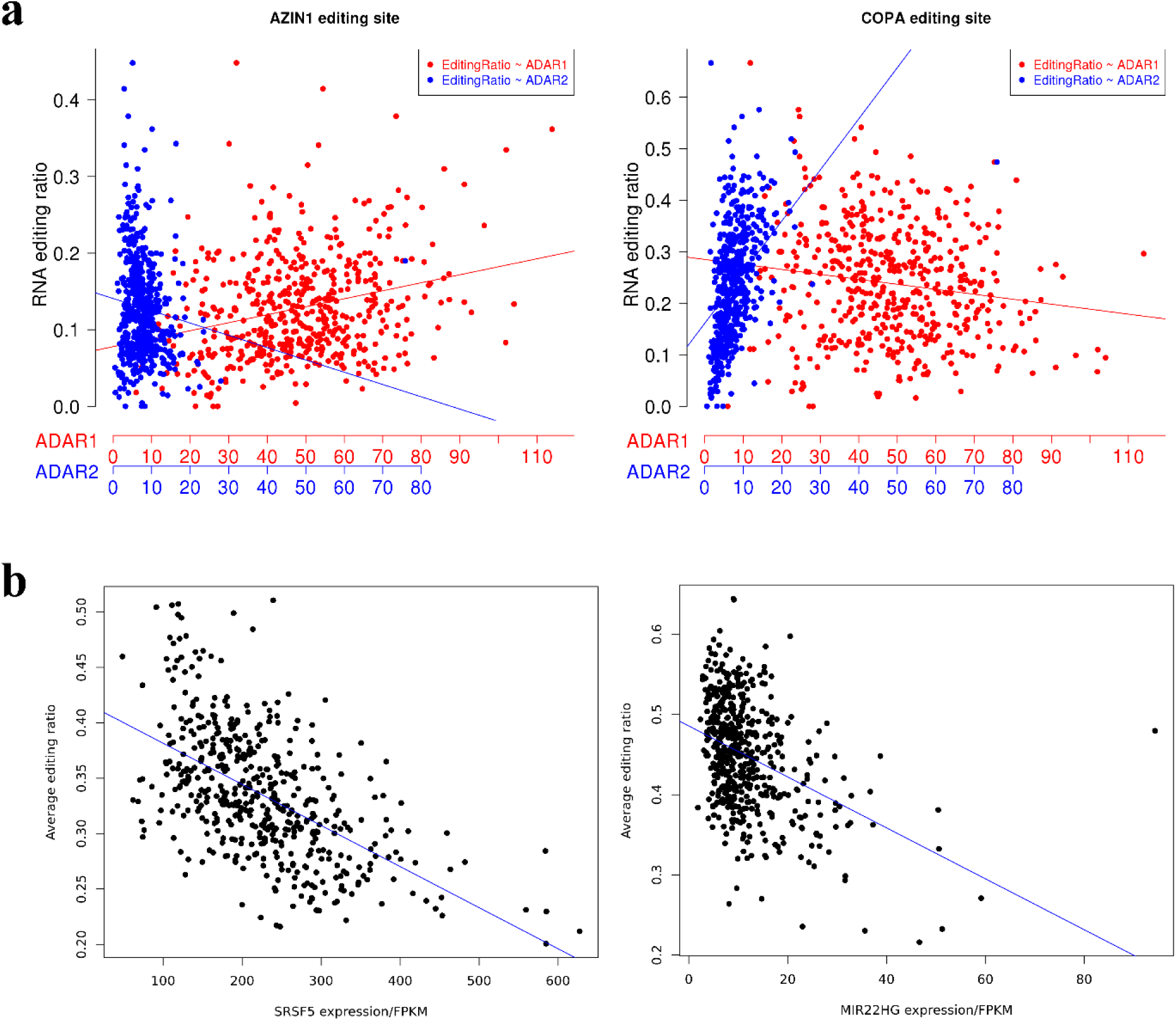
Examples of the association between RNA editing sites with ADARs and genes. (a) The editing site in AZIN1 was significantly positive associated with ADAR1 while the editing site in COPA was significantly positive associated with ADAR2, consistent with previous studies. (b) Two examples of the association between the editing sites and *SRSF5* (left), between the editing sites and *MIR22HG* (right). The average editing ratios for all the associated sites were used in plotting the figure. A linear regression line was plotted in (a) and (b) separately to show the association proximally.

### A set of novel genes were found that significantly associated with A-to-I editing sites from the second step of logistic regression

To identify the new RNA editing regulators, we did the second step of regression analysis between an RNA editing site and a gene, conditional on ADAR1, ADAR2and all the detectable interferons and their receptors (details are in method part). In the second step of regression, we only tested the editing sites that detected significantly associated with ADAR1 in the first step (the categories of ADAR1+/ADAR2#, ADAR1+/ADAR2- and ADAR1+/ADAR2+ in Table 1), which was the largest category containing ADAR significantly associated sites (Table 1). We chose the genes that significantly associated with RNA editing sites while ADAR1 was not in the second step regression as the initial set of RNA editing associated genes (the categories of ADAR1#/Gene+ and ADAR1#/Gene-in Table 3), which means these genes can better explain the editing efficiency variation across samples than ADAR1 and can be potential stronger RNA editing regulators (Model shown in Figure 1b). Most of these genes were only associated with one or two editing sites (Supplementary Figure 1), indicating the majority of the genes were either unique to specific site or artifacts. To increase confidence and gain more functional RNA editing associated genes, we only kept the genes that significantly associated with at least 20 editing sites. We further performed a permutation test and found the maximum numbers of editing sites that randomly positively associated with genes were 2, 3, 4 and 4 in KIRC, THCA, LIHC and LUAD respectively; and the maximum numbers that randomly negatively associated with genes were 3, 3, 4 and 3, much lower than the filter we used (20 editing sites), which means all RNA editing associated genes we found should not be randomly associated with RNA editing process. A total of 141, 312, 13 and 11 positive associated (PA) genes and a total of 117, 1168, 15 and 11 negative associated (NA) genes were found in KIRC, THCA, LIHC and LUAD (Table 4), respectively. All the genes are listed in Supplementary Table 2.

**Table 3.** The number of editing sites in ten categories calculated based on the association of RNA editing with ADAR1 and a new gene in KIRC, THCA, LUAD and LIHC.

|  | KIRC | THCA | LUAD | LIHC |
| --- | --- | --- | --- | --- |
| ADAR1+/Gene+ | 950 | 1040 | 1175 | 705 |
| ADAR1+/Gene# | 1154 | 1427 | 1664 | 962 |
| ADAR1+/Gene- | 1079 | 1361 | 1325 | 813 |
| ADAR1#/Gene+ | 604 | 839 | 315 | 333 |
| ADAR1#/Gene# | 780 | 982 | 932 | 740 |
| ADAR1#/Gene- | 307 | 892 | 138 | 188 |
| ADAR1-/Gene+ | 1 | 0 | 2 | 1 |
| ADAR1-/Gene# | 0 | 0 | 0 | 0 |
| ADAR1-/Gene- | 0 | 0 | 0 | 0 |
| ADAR1%/Gene% | 1175 | 1491 | 1803 | 1014 |

Notes: The signs (+, -, #, %) used in this table have the same meaning as in table1. Such as, the editing sites in the category of ADAR1+/Gene+ were significantly associated with both ADAR1 and the gene while the editing sites in the category of ADAR1#/Gene+ were only significantly associated with the gene, not ADAR1. Therefore, the genes identified using the editing sites from the category of ADAR1#/Gene+ have stronger association with the editing sites than ADAR1 while the genes identified using the editing sites from the category of ADAR1+/Gene+ have comparable association as ADAR1 with the editing sites. We selected the editing sites in the categories of ADAR1#/Gene+ and ADAR1#/Gene- for discovering editing sites associated genes.

**Table 4.** The number of PA and NA genes found in each cancer and PA and NA genes common in each two cancers. The upper corner contains the absolute number and the lower corner contains the proportion, which were calculated as the number in common divided by the smaller number of the two cancers in comparison, such as 0.455 = 5/11.

|  |  | KIRC |  | LUAD |  | LIHC |  | THCA |  |
| --- | --- | --- | --- | --- | --- | --- | --- | --- | --- |
|  |  | Num of PA genes | Num of NA genes | Num of PA genes | Num of NA genes | Num of PA genes | Num of NA genes | Num of PA genes | Num of NA genes |
| KIRC | Num of PA genes | 141 | 0 | 5 | 0 | 4 | 0 | 14 | 2 |
|  | Num of NA genes | 0 | 117 | 0 | 2 | 0 | 0 | 2 | 14 |
| LUAD | Num of PA genes | 0.455 | 0 | 11 | 0 | 0 | 0 | 1 | 0 |
|  | Num of NA genes | 0 | 0.182 | 0 | 11 | 0 | 0 | 0 | 1 |
| LIHC | Num of PA genes | 0.308 | 0 | 0 | 0 | 13 | 0 | 1 | 1 |
|  | Num of NA genes | 0 | 0 | 0 | 0 | 0 | 15 | 1 | 1 |
| THCA | Num of PA genes | 0.099 | 0.017 | 0.091 | 0 | 0.077 | 0.067 | 312 | 0 |
|  | Num of NA genes | 0.014 | 0.12 | 0 | 0.091 | 0.077 | 0.067 | 0 | 1168 |

LUAD had the smallest number of PA and NA genes although LIHC also had a small number of PA and NA genes compared to KIRC and THCA. We performed several analysis to find the potential reasons for the large variance between LUAD, LIHC and KIRC, THCA. First, we checked the sample size and sequencing depth for each cancer (Supplementary Table 1). LUAD had relative lower average sequencing depth but much higher sample size than LIHC (LUAD: 101.2 Million average aligned reads and 436 samples; LIHC: 107 Million average aligned reads and 191 samples), which indicate the nature of much smaller number of editing associated genes in LUAD and LIHC was probably not only due to the sample size and sequencing depth. Second, we further randomly extracted 191 samples (the same number as in LIHC that has the lowest sample size) from KIRC and THCA and performed the same analysis. Although the total number of PA or NA genes reduced, we found much higher number of PA genes (44 from KIRC and 186 from THCA) from both KIRC and THCA. The number of NA genes for KIRC was reduced to 9 but the number of NA for THCA was still high (782). Third, we calculated the distribution of the editing sites in the following ten categories from the second step of logistic regression: ADAR1+/Gene+, ADAR1+/Gene#, ADAR1+/Gene-, ADAR1#/Gene+, ADAR1#/Gene#, ADAR1#/Gene-, ADAR1-/Gene+, ADAR1-/Gene#, ADAR1-/Gene-, and ADAR1%/Gene% (Table 3). The meaning of the signs in each category was the same as shown in the second section in Table 1 but the threshold of significance was determined from the second step of logistic regression (details in Methods). Not surprisingly, LUAD and LIHC had much smaller number of editing sites in the categories of ADAR1#/Gene+ and ADAR1#/Gene-, which were the categories used for the PA and NA genes discovery (Table 3). However, the distribution for the ADAR1+ related categories (ADAR1+/Gene+, ADAR1+/Gene# and ADAR1+/Gene-) was different for LUAD and LIHC: LUAD was the cancer that had the largest number of editing sites in the categories of ADAR1+/Gene+ (1175), ADAR1+/Gene# (1664) and ADAR1+/Gene-(1325) across the four cancer types while LIHC had the smallest number (ADAR1+/Gene+: 705; ADAR1+/Gene#: 962; ADAR1+/Gene-: 813) (Table 3). The same tendency was found for the distribution across the categories of ADAR1+/ADAR2+, ADAR1+/ADAR2# and ADAR1+/ADAR2-: LUAD had the largest number of ADAR1 associated editing sites while LIHC had the smallest number (Table 1). The more editing sites, the higher potential of more regulators can be found. These results indicate that LUAD has potentially many regulators with regulation impact comparable to ADAR1 or less while LIHC has fewer potential regulators. Although we cannot exclude the possibility that the small number of PA and NA genes for LUAD and LIHC was caused by the sample size and sequencing depth, especially for LIHC, the analysis results indicate the nature of the LUAD and LIHC plays a role. And the apparent variation of the number for PA and NA genes across the cancer types also indicates that the regulators are more likely tissue specific.

Nevertheless, there were still genes in common between two cancer types although no genes were in common in more than two cancer types. Five PA genes were appeared in both KIRC and LUAD and four PA genes were appeared in both KIRC and LIHC, which accounted for ∼45% of the LUAD and ∼31% of the LIHC PA genes respectively (Table 3). There were fourteen common genes in KIRC and THCA for both PA and NA genes. Genes common to two cancers were listed in Supplementary Table S3. These results indicate potential common factors may be involved in the regulation RNA editing in different tissues.

We collected genes that reported to be associated with RNA editing sites from three studies^8,9,14^. Breen *et al*^8^ reported four RNA binding protein (RBP) genes that coincided with a motif shared by differential edited sites they identified in brain regions. Quinones-Valdez et al^14^ reported a total of 24 RBPs from knock down experiments that accounted for at least 10% of their tested RNA editing sites in K562 and HepG2 cells. They also concluded that the functional impact of these RBPs is likely cell type specific. Tan et al^9^ validated one gene by experiments, AIMP2, that represses the RNA editing process and reported a total of 144 PA and 147 NA genes from robust linear regression using the average ratio of all the tested editing sites in the GTEx datasets. We compared our PA and NA genes with the three reported sets of RNA editing associated genes. Two genes from THCA NA set were reported in the Breen study. Six genes from THCA NA set, one gene from THCA PA *(PABPC1*), KIRC PA (*PCBP1*) and KIRC NA (*AUH*) sets were reported in the Quinones-Valdez study. Thirty two genes from THCA NA set, three genes from THCA PA set and seven genes from KIRC PA set were reported in the Tan study. We also compared the genes (291=144 + 147) reported from the Tan study with the genes (24) reported from the Quinones-Valdez study and we found three genes were overlapped. The low overlapping rate between the two studies indicates the RNA editing regulators are tissue specific. Nevertheless, multiple genes discovered in our study were supported by the three studies indicating our method worked successfully. The results further indicate the potential RNA editing regulators are tissue/cell specific. All the genes that appeared in at least one of the three published studies were listed in Supplementary Table S4.

Interestingly, one gene in THCA NA (Figure 2b) was reported in all the three published studies discussed above: *SRSF5*, a serine/arginine (SR)-rich pre-mRNA splicing factor. SRSF5 is part of the spliceosome^15,16^ and is not only critical for pre-mRNA splicing but also involved in mRNA export from the nucleus and in translation^16,17^. Multiple studies also demonstrated that SRSF5 is involved in cancer development^18,19^. Most RNA editing events happen in the nucleus before splicing. The timing and space location of SRSF5 in the cell support that SRSF5 is very likely involved in RNA editing process.

Another interesting gene is *MIR22HG* (Figure 2b), which is an NA gene common to THCA and KIRC and also reported in the Tan study. *MIR22HG* is a RNA gene that hosts the small microRNA, miR22. The function of *MIR22HG* is not well studied but reported in multiple studies that it is a potential biomarker for several cancers^20-22^. We reported here that it may also affect RNA editing process.

### Common enriched pathways were found for both PA and NA genes across the four cancers

We performed pathway enrichment analysis using Reactome^23-25^ for the PA and NA genes in the four cancers. Although a few genes were shared in the four cancers, we identified common pathways both for PA and NA genes: Immune System for the PA genes; and Metabolism of RNA and Metabolism for NA genes (Supplementary Figures 2-9 and Supplementary Tables 5-12). The results indicate the genes may function through similar pathways across the four cancer types.

In addition to the common pathways across the four cancer types, both KIRC PA and LUAD PA genes also enriched in many other pathways, such as Cell Cycle, DNA Replication, Programmed Cell Death and Signal Transduction (Supplementary Figures 2-3 and Supplementary Tables 5-6). Similarly, commonly enriched pathways were found for the NA genes for KIRC, THCA and LIHC: Signal Transduction (Supplementary Figures 6, 8-9 and Supplementary Tables 9, 11-12); and for the NA genes for THCA, LIHC and LUAD: Metabolism of Proteins (Supplementary Figures 7-9 and Supplementary Tables 10-12). The similarity across cancers support again that common pathways likely interact with RNA editing process in cancer.

### The KIRC PA genes were significantly enriched in clinical associated genes while the KIRC NA genes were not

The enrichment of immune system genes reflects the potential association between cancer development and RNA editing process. In our previous study, we found that many RNA editing sites were differentially edited between tumor and normal samples in KIRC, and a set of the differential edited sites (80) in KIRC were significantly associated with patient survival^11^. This evidence suggests that RNA editing sites impact on cancer development. Here we tested whether the candidate genes were also associated with KIRC patient survival. The Cox proportional hazards model was used for the analysis where gene expression, gender and race were covariates. We used the same thresholds to select the genes as we did in the second step of logistic regression: the average expression of the gene > 5 (FPKM) in at least 50 tumor samples. In total we tested 43823 genes, of which 10902 (∼25%) were significantly associated with patient survival (at threshold of FDR 5%). Of the 141 PA genes in KIRC, 75 (∼53%) were significantly associated with patient survival; while 22 (∼19%) of the 117 NA genes were significantly associated with patient survival. Compared with the total genes tested (48455), PA genes were enriched in survival associated genes (Chi squared test; p value is ∼1.88e-14) while the NA genes were toward to depletion of survival associated genes (Chi squared test; p value is ∼0.16). The significant difference between PA and NA genes indicates that the PA genes, which enriched in Immune System genes, seems playing more important roles in cancer development than the NA genes.

## Discussion

In this work, we developed a two-step logistic regression model to identify RNA editing associated genes in four cancer types. The first step was to identify the dominant enzyme that catalyzed the RNA editing sites, and we found that ADAR1 was the enzyme that associated with the majority of the editing sites in all four cancers. In the second step we identified the genes that had potentially stronger regulation effect on RNA editing than ADAR1. Using this approach, we identified a set of PA and NA genes in each cancer that very likely involved in regulation of RNA editing process. The number of PA and NA genes varied largely across the four cancer types: THCA had the largest number of PA (312) and NA (1168) genes while LUAD had the smallest number of PA (11) and NA (11) genes. Most of the genes were unique to each cancer but some overlapped (∼45% (5) of LUAD PA and ∼31% (4) LIHC PA genes were common to KIRC PA genes). Interestingly, the PA genes were enriched in Immune System pathway and the NA genes was enriched in Metabolism of RNA and Metabolism pathways, indicating the genes may function through common pathways across cancer. More importantly, we found that the PA genes in KIRC were enriched in survival-associated genes, while the NA genes were depleted of survival-associated genes.

There are multiple advantages to use the logistic regression in two steps. First, we quantified the efficiency of RNA editing not simply based on the RNA editing ratio (the number of edited reads/the number of total reads), but estimated the expected editing ratio based on the number of edited and un-edited reads at each site under the assumption that the expected editing ratio is the successful rate of observing an event. This method can take the sequencing depth into the consideration, thus, can quantify the editing efficiency in a much more accurate way. Second, the first step was able to group the RNA editing sites into different categories based on the association with ADARs. Thus we can test the method based on the known pairs of editing sites and the enzymes. Third, the first step classified the editing sites into 10 categories which make the following analysis easier and more focused, especially identifying that ADAR1 was the enzyme that associated with the majority of editing sites. Therefore the editing sites associated with ADAR1 were taken into the second step analysis. Fourth, the second step took ADAR1 as a covariate in the regression to identify the candidate genes that have strong regulation effect by comparing the effect of ADAR1 with a new gene, and selected the genes that had stronger effect than ADAR1. This was the reason we selected the genes in the categories of ADAR1#Gene+ and ADAR1#Gene-rather than ADAR1+Gene+ and ADAR1+Gene-. Fifth, in the second step we also took the detectable interferons and their receptors as covariates in the analysis to account for potential artificial association between editing sites and the candidate genes caused by interferons. The advantages of our approach create a promising method to identify confident factors involved in RNA editing process. The idea of our algorithm can be used in similar studies, such as other functional processes where known factors can serve as controls like ADARs.

In this study, the highest number of editing sites that one candidate gene can control was 495 which was smaller than the number of sites tested (839), suggesting that different sets of RNA editing sites have potentially specific regulators (Supplementary Figure 1). Moreover, most genes were associated with one editing site although they were filtered in our analysis (Supplementary Figure 1). After breaking the association of editing sites with genes in the permutation test, many of the genes were not associated with any editing site, indicating some genes, even associated with one editing site, can still be true candidate genes. Interestingly we also found that some editing sites were associated with only one candidate gene, and some editing sites can be associated with more than 2000 candidate genes (Supplementary Figure 10). Previous studies showed that different editing sites have different effects on cancer, either repress or promote cancer progress at different levels, which not only demonstrate the complexity of RNA editing effects, but also the site specific effect^26^. We hypothesize that the site-specific effects may be caused by site specific regulators.

Although our method took ADAR1 as a control gene, the PA and NA genes we identified may function through interacting with ADAR1 or not. Our model only indicates that these PA and NA genes were the factors that explained a higher variance than ADAR1 on editing efficiency across samples for the associated editing sites. Whether these genes directly interact with ADAR1 or indirectly interact with ADAR1 or not interact with ADAR1 at all need further analysis and experiments to reveal.

## Methods

### Association analysis between RNA editing sites and the expression of ADAR1 and ADAR2 (the first step of logistic regression)

We used a logistic regression model to find the corresponding enzyme for each editing site. The model was shown below:

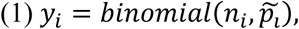

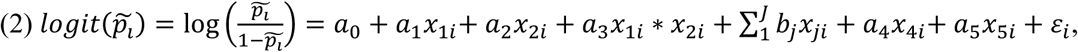

*y*_*i*_ is the number of edited reads at one RNA editing site. *n*_*i*_ is the number of total reads at one RNA editing site 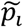 is the expected RNA editing ratio, which was estimated as the probability of success in the binomial distribution (model 1). *x*_1*i*_ is the expression of ADAR1 and *x*_2*i*_ is the expression of ADAR2. *x*_1*i*_ * *x*_2*i*_ is the interaction term between ADAR1 and ADAR2. J is the total number of interferons and their receptors. *x*_*ji*_ is the expression of the interferon or its receptor. *x*_4*i*_is the gender. *x*_5*i*_ is the race. *i* represents the *i* ^th^ sample. *j* represents the *j* th interferon or its receptor. The estimated editing ratio was logit transformed to convert the ratio from 0 to 1 to -∞ to +∞. And then the transformed ratio was used to fit a linear regression (model 2). This approach took the sequencing coverage at each site into consideration. Therefore, the model can handle overdispersion data. 3,800 RNA editing sites were tested in the analysis. But we filtered the sites if they were not detectable in at least 50 samples. To determine the significantly associated enzyme, we used a threshold of p value at 1.3e-05, which was proximal to a q value of 0.05 after Bonferroni correction (∼0.05/3800). If the coefficiency was higher than 0, we classified the editing site as a positive associated site with the enzyme (such as ADAR1+), otherwise, the editing site was classified into the negative associated groups (such as ADAR1-). If the p value was higher than 1.3e-03, the editing site was classified as not significantly associated with the enzyme, such as the categories of ADAR1# and ADAR2#. The sites in the categories of ADAR1# and ADAR2# may not be absolutely associated with ADARs since here we used a stringent criterion to identify the sites that had very strong association with ADARs. Any site that cannot be grouped into any of the nine categories mentioned in the main text was classified as an undefined group, marked as ADAR1%/ADAR2%. The results were summarized in Table 1.

### Association analysis between RNA editing sites and gene expressions (the second step of logistic regression)

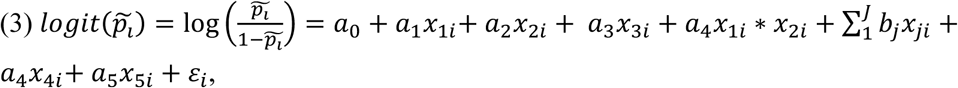

The model 3 was similar to model 2 except we added a covariate, *x*_3*i*_, which is the expression of a gene except, ADAR1, ADAR2, the interferons and their receptors. The expected RNA editing ratio, 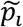 was estimated as described in the first section (model 1). The expression of ADAR1 in the model was used as a control gene. We selected the genes as candidates if they had stronger association with editing sties than ADAR1. Thus, we selected the genes that were significant associated with editing sites while the ADAR1 was not, which indicates the candidates can explain the variance of the editing sites across samples more than ADAR1. A total number of 55401 genes were initially analyzed, but we filtered the genes with an average expression of less than 5 FPKM and also the genes expressed in less than 50 samples. We used a threshold of p value of 2.3e-10 to select the significantly associated genes, which was proximal to a q value of 0.05 after Bonferroni multiple testing correlation (0.05/55401/3800). If the coefficiency was higher than 0, we classified the editing site as a positive associated site with the gene, otherwise, the editing site was classified into negative associated groups (such as Gene+ or Gene-). If the p value was higher than 2.3e-07, the editing site was classified as not significantly associated with the gene, such as the related categories of Gene# and Gene#. Any site that cannot be grouped into any of the three categories (Gene+, Gene- and Gene#) was classified as an undefined group, marked as Gene%. The number of editing sites analyzed in each category were summarized in Table 3.

### Permutation test

We did the permutation for each site and each gene. In the permutation test, we kept the relationship among all the editing sites and among all the gene expressions, but only interrupted the connection between the RNA editing site and the gene expression in each regression. Each test was permutated 100 times and then we fit the same models (mode 3 and 4) as described above. Then the number of RNA editing sites each gene significantly associated with was summarized in the main text. No gene was associated with more than four editing sites in all the four cancers.

### Pathway enrichment analysis

We used Rectome to do the analysis for the PA and NA genes obtained from the four cancers. We uploaded the candidate genes to rectome (https://reactome.org/) and performed the analysis using all the genes in Homo sapiens as the reference list. The default settings were used for all the analysis.

### Survival analysis using Cox proportional hazards model

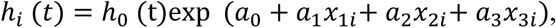

where *h*_*i*_ (*t*) is the hazard function of the *i* th individual at time *t*, and *h*_0_ (t) is the baseline hazard. *x*_1*i*_ is gene expression. *x*_2*i*_ is the gender. *x*_3*i*_ is the race. *i* represents the *i* ^th^ sample. The calculation was conducted using the function of coxph from the package of survival. Genes with an average expression of less than 5 FPKM and also expressed in less than 50 samples were filtered out in the survival analysis.

## Author Contributions Statement

TG conceived and designed the project, developed the method, and performed the analysis. AF, MB and XZ performed partial analysis. TG drafted the manuscript and all authors edited the manuscript.

## Conflict of Interest Statement

The authors declare no competing interests.

