## Supplemental Figures for "Systematic identification of A-to-I editing associated regulators from multiple human cancers"

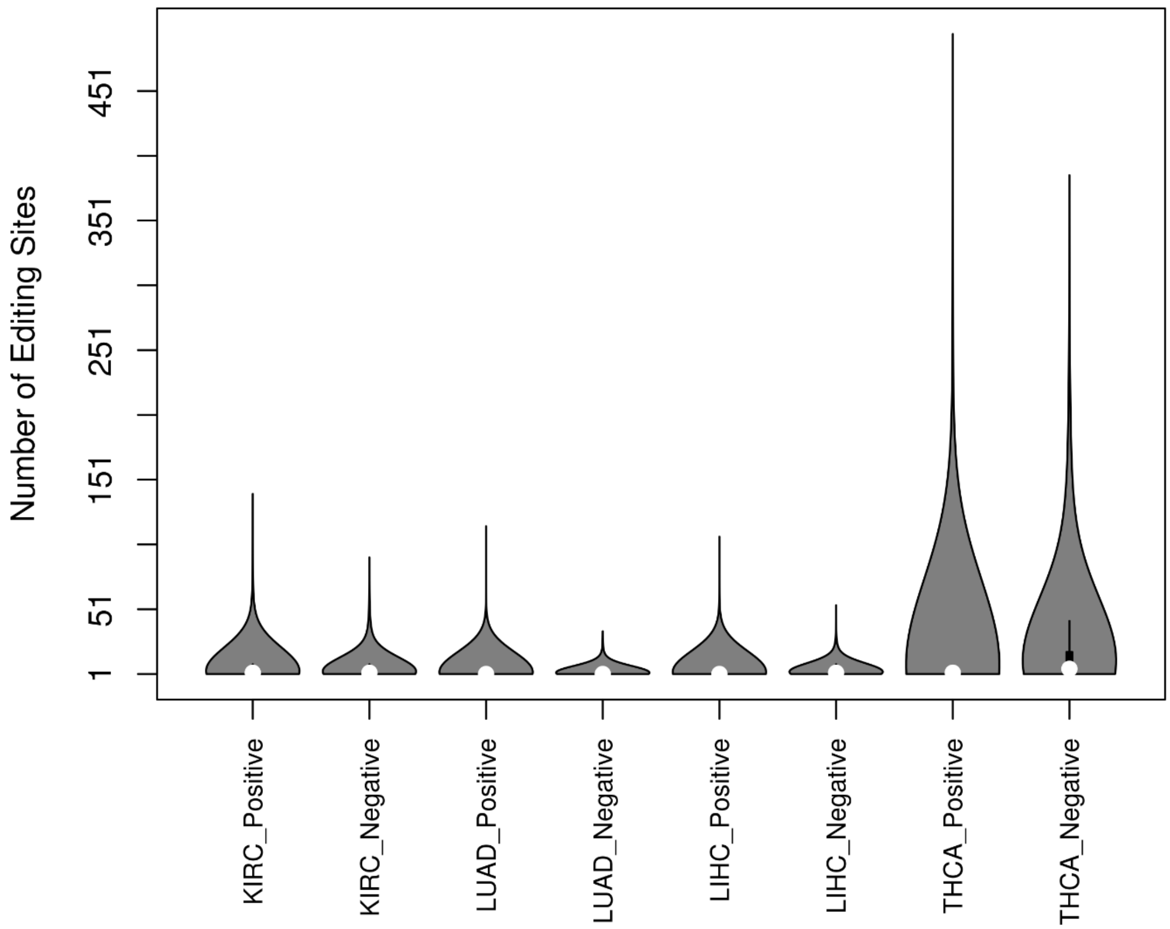


Supplementary Figure 1, Distribution of the number of editing sites that each gene associated with in the four cancers. Most candidate genes control one or two sites, but they can regulate as many as more than 450 RNA editing sites.

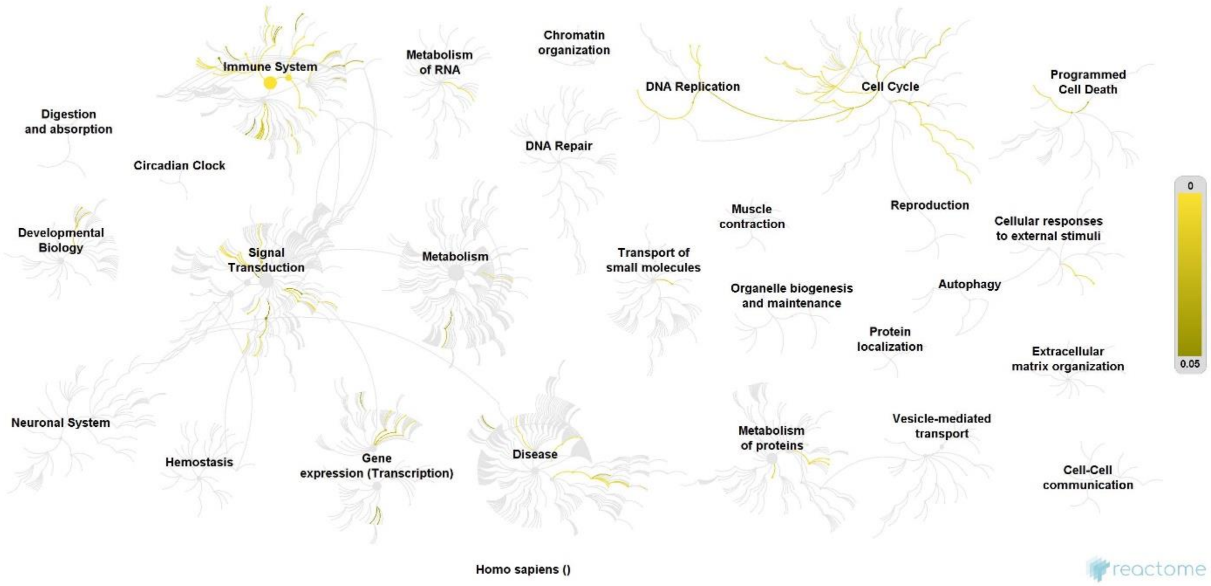


Supplementary Figure 2, Enriched pathways for the KIRC PA genes. The yellow color marks the P value from 0.05 to 0.

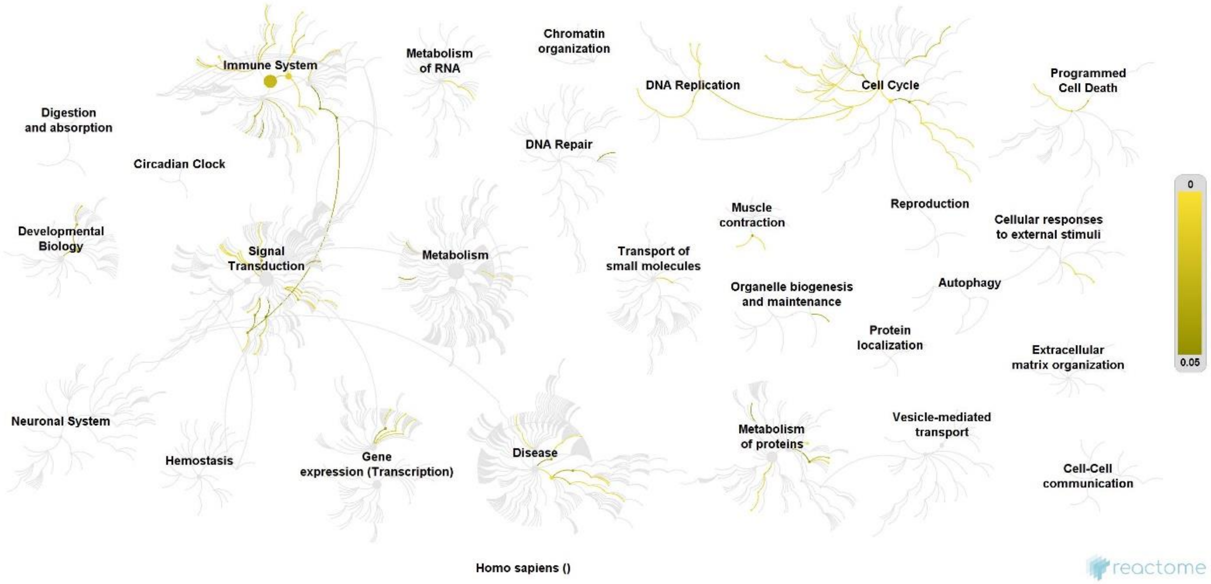


Supplementary Figure 3, Enriched pathways for the LUAD PA genes. The yellow color marks the P value from 0.05 to 0.

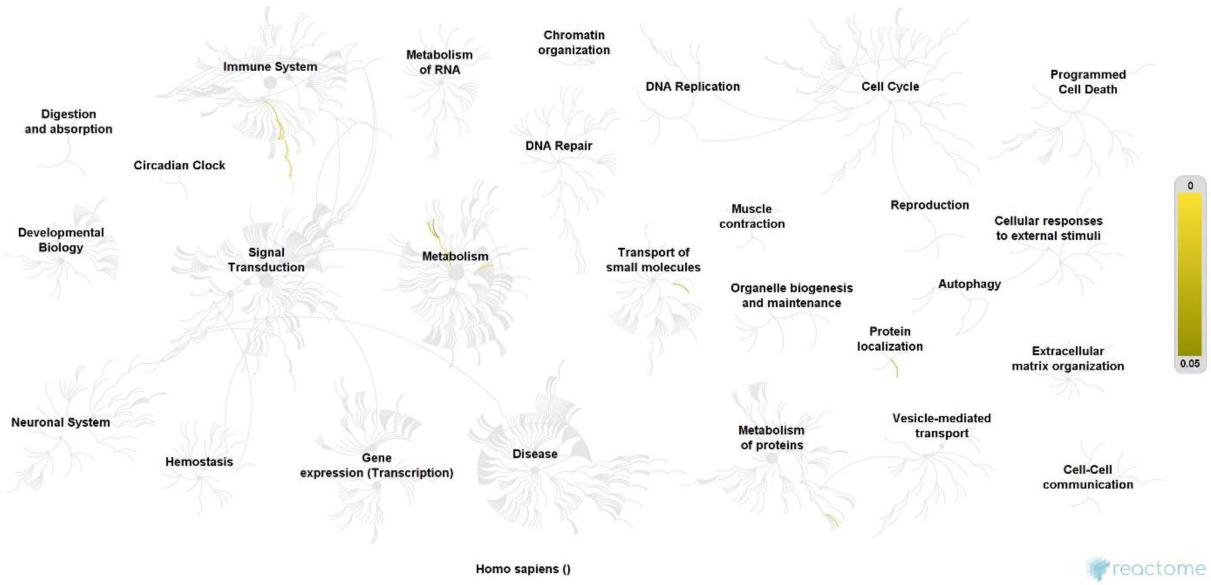


Supplementary Figure 4, Enriched pathways for the LIHC PA genes. The yellow color marks the P value from 0.05 to 0.

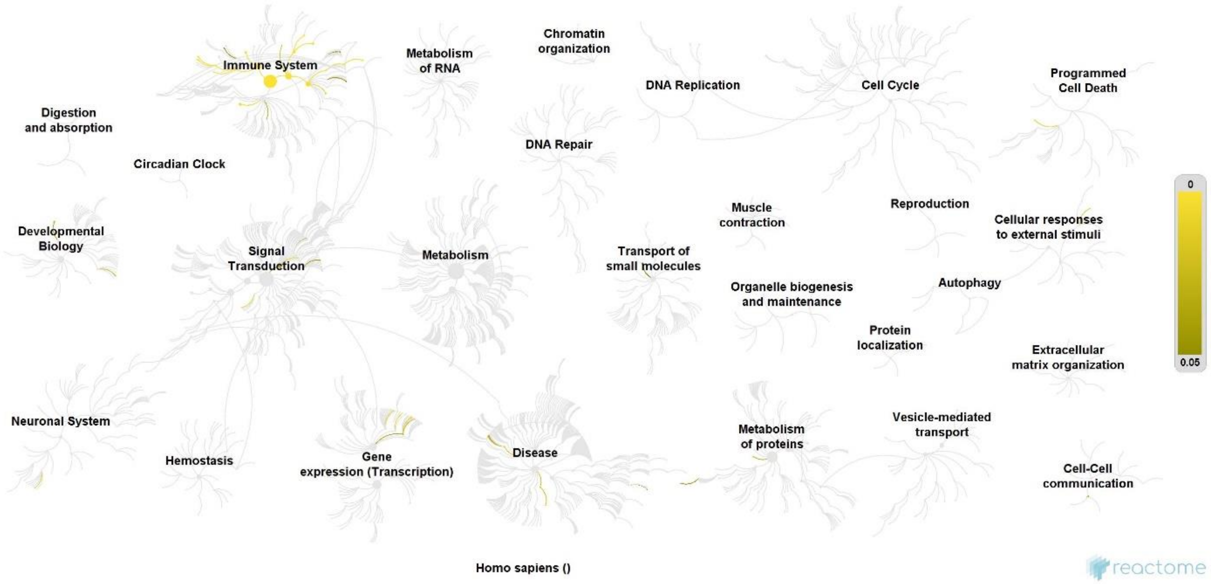


Supplementary Figure 5, Enriched pathways for the THCA PA genes. The yellow color marks the P value from 0.05 to 0.

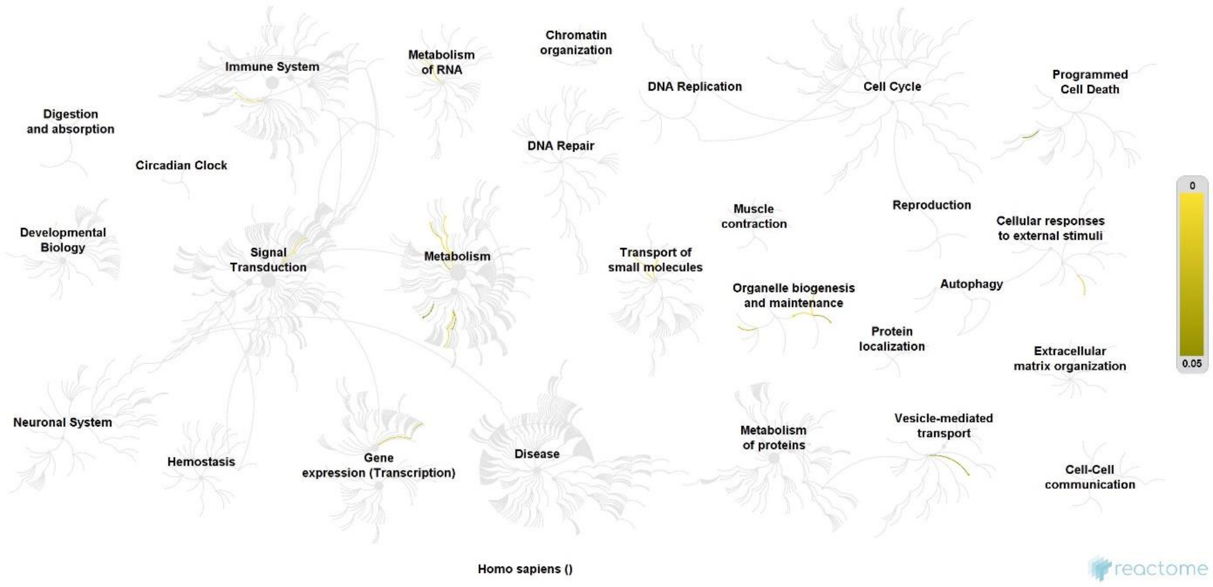


Supplementary Figure 6, Enriched pathways for the KIRC NA genes. The yellow color marks the P value from 0.05 to 0.

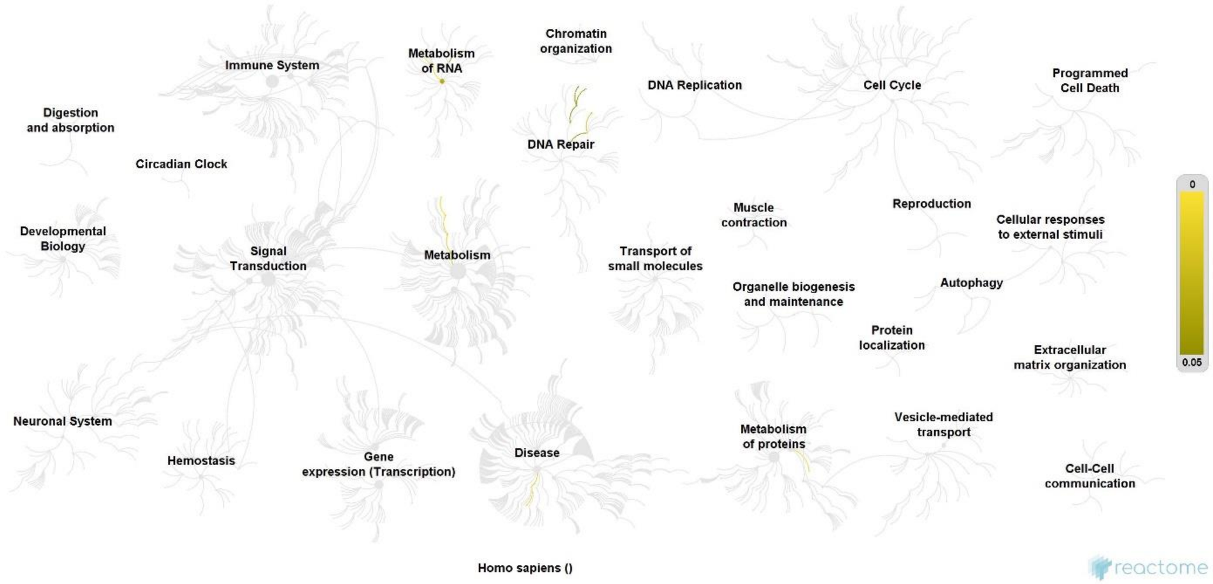


Supplementary Figure 7, Enriched pathways for the LUAD NA genes. The yellow color marks the P value from 0.05 to 0.

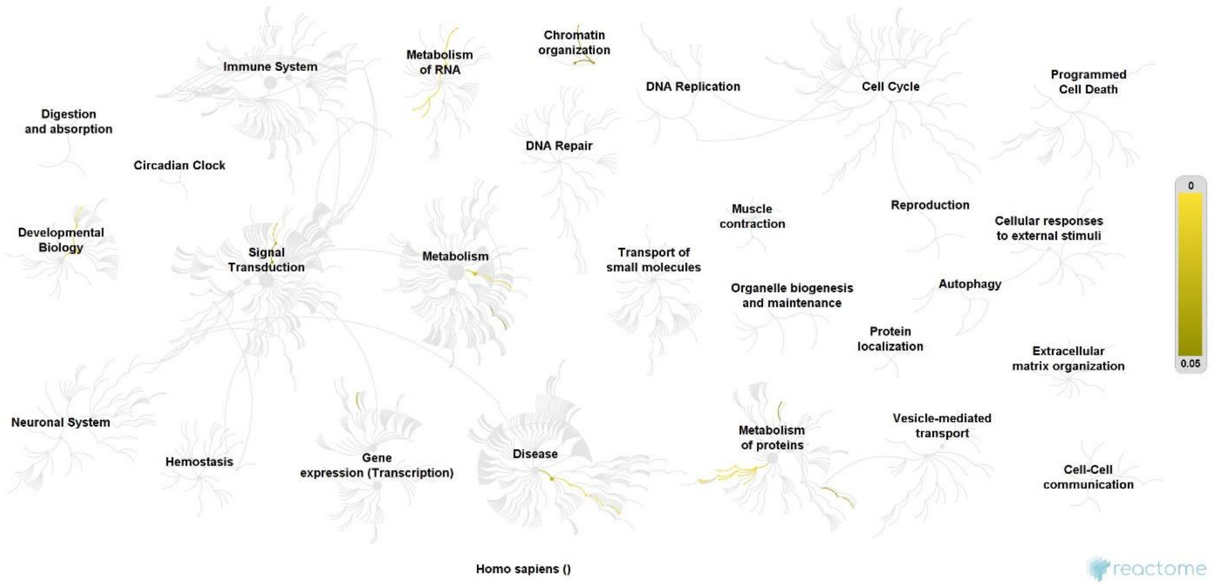


Supplementary Figure 8, Enriched pathways for the LIHC NA genes. The yellow color marks the P value from 0.05 to 0.

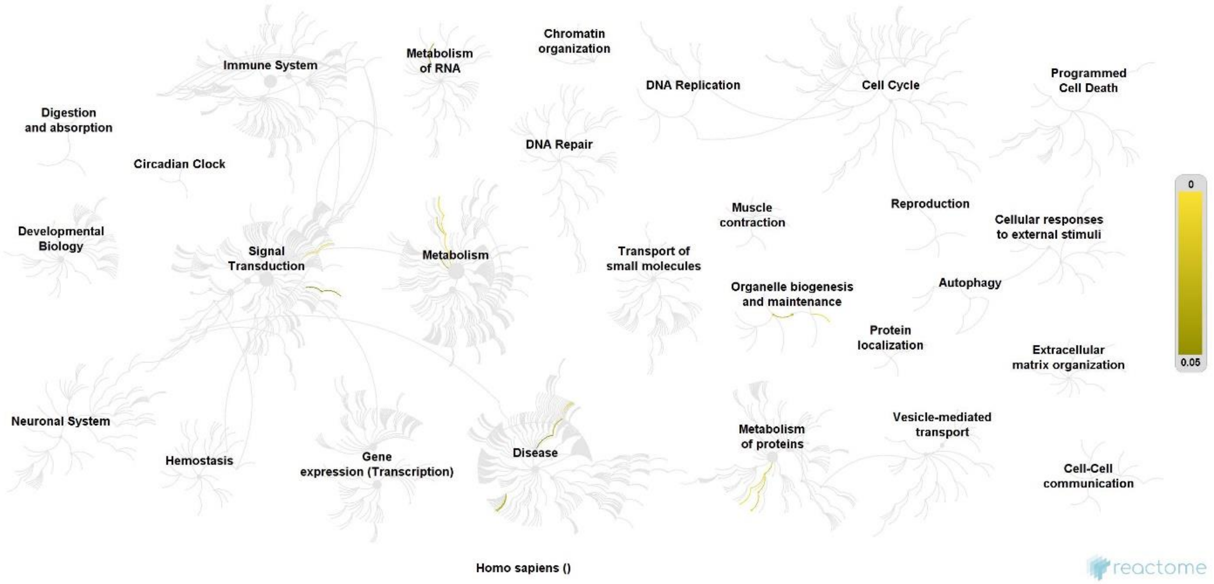


Supplementary Figure 9, Enriched pathways for the THCA NA genes. The yellow color marks the P value from 0.05 to 0.

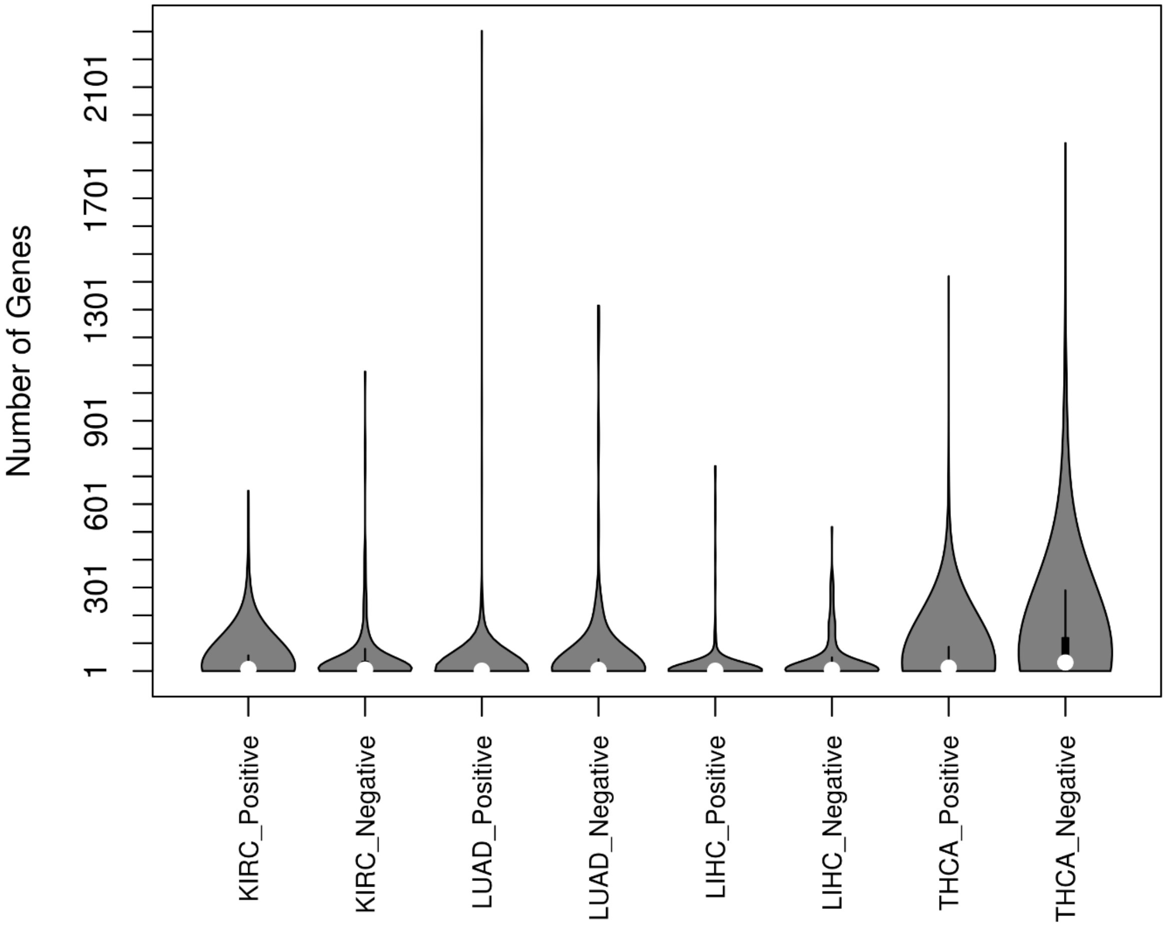


Supplementary Figure 10, Distribution of the number of genes that each editing site associated with in the four cancers.
